# Type 1 interferon supports B cell responses to polysaccharide antigens but is not required for MPL/TDCM adjuvant effects on innate B cells

**DOI:** 10.1101/2021.10.30.466603

**Authors:** M. Ariel Spurrier, Jamie E. Jennings-Gee, Karen M. Haas

## Abstract

We previously described monophosphoryl lipid A (MPL) and synthetic cord factor, trehalose-6,6’-dicorynomycolate (TDCM) significantly increases antibody (Ab) responses to T cell independent type 2 antigens (TI-2 Ags) in a manner dependent on B cell-intrinsic TLR4 expression as well as MyD88 and TRIF adapter proteins. Given the requirement for TRIF in optimal MPL/TDCM adjuvant effects and the capacity of MPL to drive type I IFN production, we aimed to investigate the extent to which adjuvant effects on TI-2 Ab responses depend on type I IFN receptor (IFNAR) signaling. We found IFNAR^−/−^ mice had impaired early TI-2 Ag-induced B cell activation and expansion and that B cell-intrinsic type I IFN signaling on B cells was essential for normal antibody responses to TI-2 Ags, including haptenated Ficoll and the pneumococcal vaccine, Pneumovax23. However, MPL/TDCM significantly increased TI-2 IgM and IgG responses in IFNAR^−/−^ mice. MPL/TDCM enhanced TI-2 Ab production primarily by activating innate B cells (B-1b and splenic CD23^−^ B cells) as opposed to CD23^+^ enriched follicular B cells. In summary, our study highlights an important role for type I IFN in supporting early B cell responses to TI-2 Ags through B cell-expressed IFNAR, but nonetheless demonstrates an MPL/TDCM adjuvant significantly increases TI-2 Ab responses independently of type I IFN signaling and does so by predominantly supporting increased polysaccharide-specific Ab production by innate B cell populations.

**Key points:**

- B cell-intrinsic IFNAR expression promotes TI-2 Ab responses.
- MPL/TDCM adjuvant effects are independent of type 1 IFN.
- MPL/TDCM promotes TI-2 Ab responses by innate B cells.

## Background

Pneumococcal vaccines consist of *S. pneumoniae*-derived capsular polysaccharides (PPS) which behave as T cell-independent type 2 antigens (TI-2 Ags). Pneumovax^®^23, composed of 23 types of PPS, provides protection against invasive pneumococcal disease in adults for up to 10 years, with an estimated efficacy of 60-70% (1). Adjuvants for native polysaccharide vaccines have not been used in the clinic. Alum does not boost primary Ab responses to TI-2 Ags (2) and poorly boosts responses to PPS-conjugates (3). Earlier work with Ribi adjuvant, consisting of *Salmonella typhimurium* monophosphoryl lipid A (MPL) and *Mycobacterium* cord factor in squalene elicited increased primary PPS-specific Ab responses in mice (4, 5), but is not suitable for use in humans due to its toxicity. We recently demonstrated low-toxicity *Salmonella minnesota* MPL and synthetic cord factor analog trehalose dicorynomycolate (TDCM) emulsified in squalene significantly increased Ag-specific IgM and IgG levels in response to PPS and haptenated-Ficoll (6). The mechanisms by which this adjuvant carries out its effects have not been fully elucidated but is important for informed design of adjuvants for use with polysaccharide Ags in humans.

We previously reported MPL/TDCM induced type I IFN production in vivo (7), likely through the effects of MPL on TLR4-TRIF activation of type I IFN production (8). Importantly, a previous study demonstrated the TLR3 agonist, poly(I:C), increased Ab responses to NP-Ficoll via a mechanism dependent upon IFNAR signaling and follicular B cells (9). To further understand the mechanisms by which MPL/TDCM increases Ab responses to TI-2 Ags, we investigated the importance of type I IFN and B cell subsets in adjuvant effects. Herein, we demonstrate that despite finding a clear role for B cell-intrinsic IFNAR signaling in supporting B cell responses to TI-2 Ags, MPL/TDCM adjuvant effects are not dependent on type I IFN signals. Further, we demonstrate MPL/TDCM primarily elicits its effects by stimulating increased Ab production by innate B cell subsets (ie., B-1b and splenic CD23^−^ populations), as opposed to splenic CD23^+^ cells which largely represent follicular B cells. These findings highlight the importance of type I IFN in TI-2 Ab responses and reveal distinct pathways that may be leveraged for improving TI-2 Ab responses.

## Methods

### Mice

Wild-type (WT), mumt (Ighm^tm1Cgn^), CD19^Cre^(B6.129P2(C)-Cd19^tm1(cre)Cgn^/J) and B1-8hi IgH knock-in (V_H_B1-8 Tg) mice were on a C57BL/6 background (Jackson Laboratories). Ifnar1-deficient (IFNAR^−/−^) and floxed IFNAR^fl/fl^ (Ifnar1^tm1Uka^) mice were as previously described (7). Studies used age- and sex-matched mice housed in an SPF facility and were approved by Wake Forest School of Medicine’s Animal Use Committee.

### Immunizations and ELISAs

Mice were immunized with 10 μg TNP65-Ficoll, 1 μg NP_40_-Ficoll, or the equivalent of ~0.1 μg each PPS within Pneumovax23 (Merck). Adjuvant containing 20 μg *Salmonella minnesota* MPL, 20 μg trehalose-6,6‘-dicorynomycolate (TDCM) in 0.5% squalene/0.05% Tween-80 or 2% squalene/0.2% Tween-80 (Sigma) for intraperitoneal (i.p.) and intramuscular (i.m.) injections, respectively, was mixed with Ag prior to injection. ELISAs were as previously described (6, 10, 11). TNP- and PPS-specific Ig levels were estimated using a standard curve generated using anti-mouse Ig (H+L) coated wells in conjunction with mouse IgM and IgG isotype standards (Southen Biotechnology Associates). NP-specific Ig concentrations were estimated using NP-specific IgM and IgG standard curves as previously described (11).

### Flow cytometry

Peritoneal cells were harvested using 10 ml of DPBS to lavage the peritoneal cavity. Splenocytes and peritoneal cells were resuspended in staining buffer (PBS containing 2% newborn calf serum) and pre-incubated with 0.5 μg/ml FcBlock (eBioscience) and stained with fluorochrome-conjugated mAbs (Biolegend, eBioscience, and BD Biosciences): CD19 (1D3), CD11b (M1/70), CD5 (53-7.3), CD23 (B3B4), CD21/35 (7E9), CD138 (281-2), CD86 (GL1), CD19 (1D3), CD11b (M1/70), CD45R/B220 (RA3-6B2), rat anti-mouse IgG (pooled rat anti-mouse IgG1, IgG2b, IgG2a, IgG3; Southern Biotechnology Associates), and Live/Dead fixable aqua stain. After staining, samples were washed with DPBS + 2% FCS and fixed in 1.5% buffered formaldehyde. For TNP-specific B cell staining, cells were incubated with TNP(65)-FL-AECM-Ficoll (20μg/ml; Biosearch Technologies Inc.) prior to performing extracellular staining. Ki-67 staining (solA15) was performed according to manufacturer’s instructions (eBioscience™ Foxp3/ Transcription Factor Staining Buffer Set). Cells were analyzed using a BD LSR Fortessa X20 (Becton Dickinson).

### Bone marrow chimeras

WT recipient mice were lethally irradiated (950 rad) and reconstituted i.v. with 10^7^ total bone marrow cells consisting of muMT bone marrow mixed with either WT or IFNAR^−/−^ bone marrow in a 90:10 ratio as previously described (12). Recipient mice were maintained on Septra 1 week prior to irradiation and 2 weeks afterwards. Mice were rested for 4 weeks prior to immunization.

### Adoptive transfer experiments

Naïve CD43^−^ splenic and peritoneal B cells were purified from V_H_B1-8 Tg mice using CD43 bead depletion (Dynal). CD23^+^ and CD23^−^ B cells were further purified using Miltenyi beads. B-1b cells were further enriched using CD11b Miltenyi beads. V_H_B1-8 Tg B cells were transferred i.v. into CD45.2^+^ mice. Naïve spleen B cells were purified from WT mice in a similar manner and WT peritoneal B cells were obtained using EasySep pan B cell isolation (StemCell) combined with biotinylated F4/80. A second purification step was used for B-1b cell enrichment using biotinylated Abs against CD5, F4/80, and GR1 in conjunction with streptavidin Dynabeads (>90% CD5^−^CD19^+^ B cell purity; 70% CD11b^+^). Recipient muMT mice were reconstituted with cells i.v.

### Statistical analyses

Data are shown as means ± SEM. Differences between sample means were assessed using Student’s t test.

## Results

### Type I IFN signaling is required for optimal Ab responses to TI-2 Ags

Before investigating the role of type I IFN signaling in MPL/TDCM adjuvant effects, we first investigated whether type I IFN played a role in regulating Ab responses to TI-2 Ags. As shown in Fig. 1A, mice lacking IFNAR1 and thus functional IFNAR (IFNAR^−/−^) exhibited a significant reduction (~30%) in TNP-specific IgM and IgG levels 7 days post TNP-Ficoll immunization, although responses approximated WT levels at later time points. However, in response to Pneumovax23 immunization, IFNAR^−/−^ mice produced significantly less (50%) IgM and IgG to serotype 3 polysaccharide (PPS3) found within the vaccine as well as significantly less IgM and IgG reactive with total Pneumovax23, and these differences were sustained (Fig. 1B-C). Thus, IFNAR supports optimal TI-2 Ab responses, and is particularly important for normal IgM and IgG responses to pneumococcal polysaccharides.

**Figure 1.**
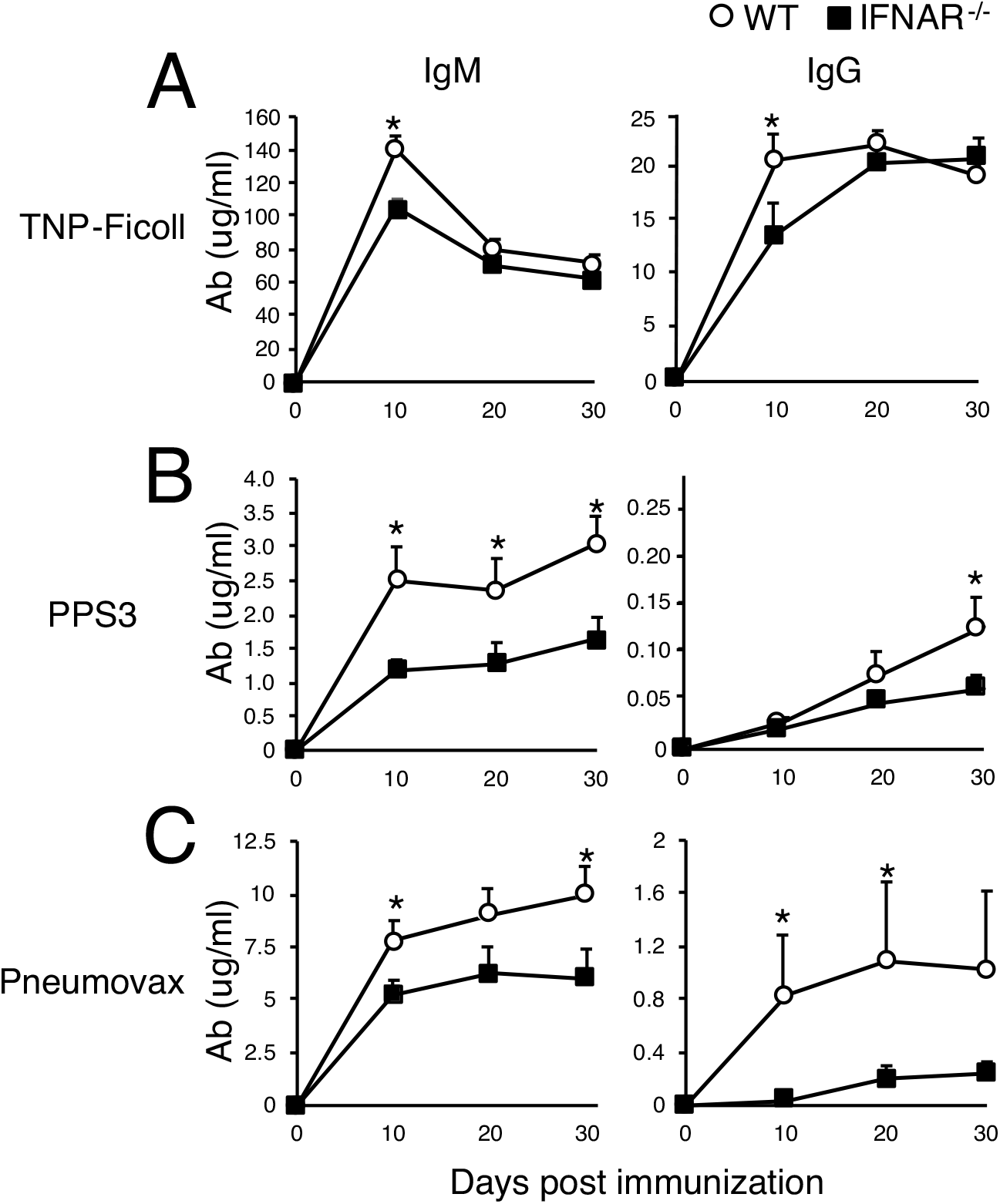
Type I IFN receptor signaling is required for optimal Ab responses to TI-2 Ags. A-C) WT and IFNAR1^−/−^ mice were immunized with 25 μg TNP65-Ficoll or Pneumovax23 containing 1 μg each PPS. Serum IgM and IgG levels reactive with TNP-BSA (A), PPS3 (B), and whole Pneumovax23 (C) were determined by ELISA (n≥5 mice/group). Asterisks (*) indicate significant differences between Ab levels in WT and IFNAR^−/−^ mice. Results are representative of those obtained in 4 independent experiments.

### B cell subset development is normal in IFNAR^−/−^ mice

As alterations in innate B cell subsets can lead to defective TI-2 Ab responses, we assessed B cell subsets in IFNAR^−/−^ mice. As shown in Fig. 2, we found total B cell frequencies and numbers in spleen and peritoneal cavity were similar between WT and IFNAR^−/−^ mice. Furthermore, we did not find differences in B cell subsets. Peritoneal B-2 cell and spleen follicular B cell frequencies and numbers were similar in WT and IFNAR^−/−^ mice as were innate peritoneal B-1a and B-1b and splenic marginal zone B cell frequencies and numbers (Fig. 2). Thus, B-1 and B-2 B cell subset development and maintenance is normal in IFNAR^−/−^ mice and does not explain impaired TI-2 Ab responses in IFNAR^−/−^ mice.

**Figure 2.**
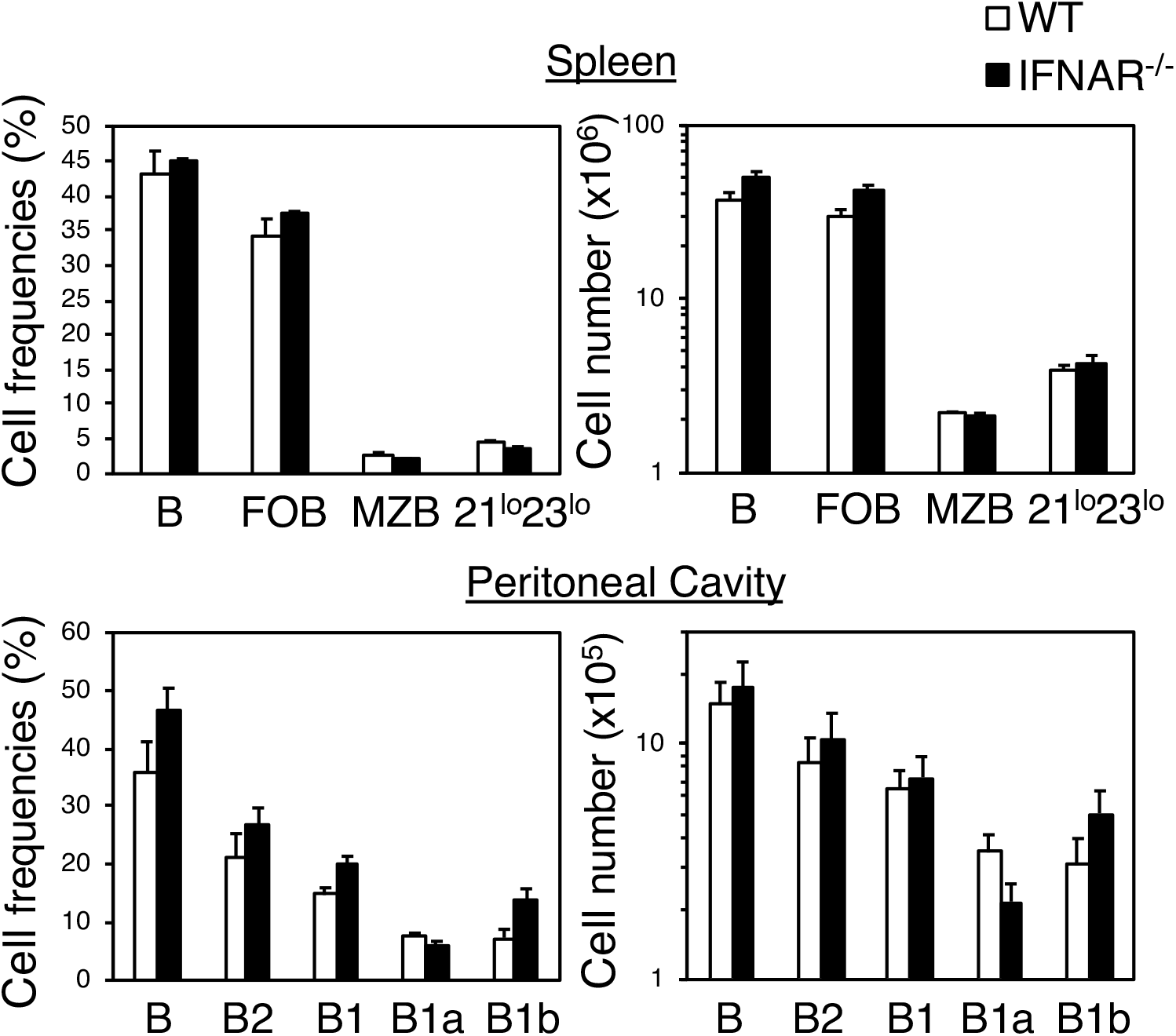
Normal B cell subset distribution in IFNAR^−/−^ mice. Spleen and peritoneal B cell subsets were analyzed in naïve WT and IFNAR1^−/−^ mice. The mean frequency of total spleen B (CD19^+^), follicular B (FOB:CD21^int^CD23^+^), marginal zone B (MZB: CD21^hi^CD23^−^) and CD21^lo^CD23^lo^ cells among total leukocytes and numbers are indicated in the top panels. The frequency of total peritoneal B (CD19^+^), B-2 (CD11b^−^), B1 (CD11b^+^), B1a (CD11b^+^CD5^+^), and B1b (CD11b^+^CD5^−^) cells among total leukocytes and numbers are indicated in the bottom panels (n=5 mice/group).

### Type I IFN signaling on B cells is required for optimal Ab responses to TI-2 Ags

IFNAR expression is ubiquitous and hence, altered TI-2 Ab responses in IFNAR^−/−^ mice could be due to lack of IFNAR on one or more cell types. We investigated the importance of B cell-expressed IFNAR in TI-2 Ab responses by crossing IFNAR1^fl/fl^ mice with CD19-Cre transgenic mice to generate mice lacking IFNAR only on CD19-expressing B cells (CD19-Cre^+/-^IFNAR^fl/fl^) or expressing heterozygous IFNAR expression on B cells (CD19-Cre^+/-^IFNAR^fl/+^). As shown in Figure 3, CD19-Cre^+/-^ mice produced lower Ab responses to TNP-Ficoll and PPS relative to WT mice and were therefore used to draw comparisons with CD19-Cre^+/-^IFNAR^fl/fl^ mice. In response to TNP-Ficoll, CD19-Cre^+/-^IFNAR^fl/fl^ mice produced significantly less TNP-specific IgG relative to CD19-Cre^+/-^ and CD19-Cre^+/-^IFNAR^fl/+^ mice (Fig. 3A). In response to Pneumovax23, CD19-Cre^+/-^IFNAR^fl/fl^ mice produced significantly less PPS3-specific IgM, and PPS3- and Pneumovax23-specific IgG levels were 50% decreased relative to CD19-Cre^+/-^ mice (Fig. 3B-C). Interestingly, CD19-Cre^+/-^IFNAR^fl/+^ mice expressing heterozygous IFNAR levels on B cells produced TI-2 Ab levels similar to CD19-Cre^+/-^ mice, suggesting heterozygous IFNAR expression on B cells is sufficient to support B cell responses to TI-2 Ags, whereas selective deficiency of IFNAR on B cells results in impaired Ab responses to TI-2 Ags.

**Figure 3.**
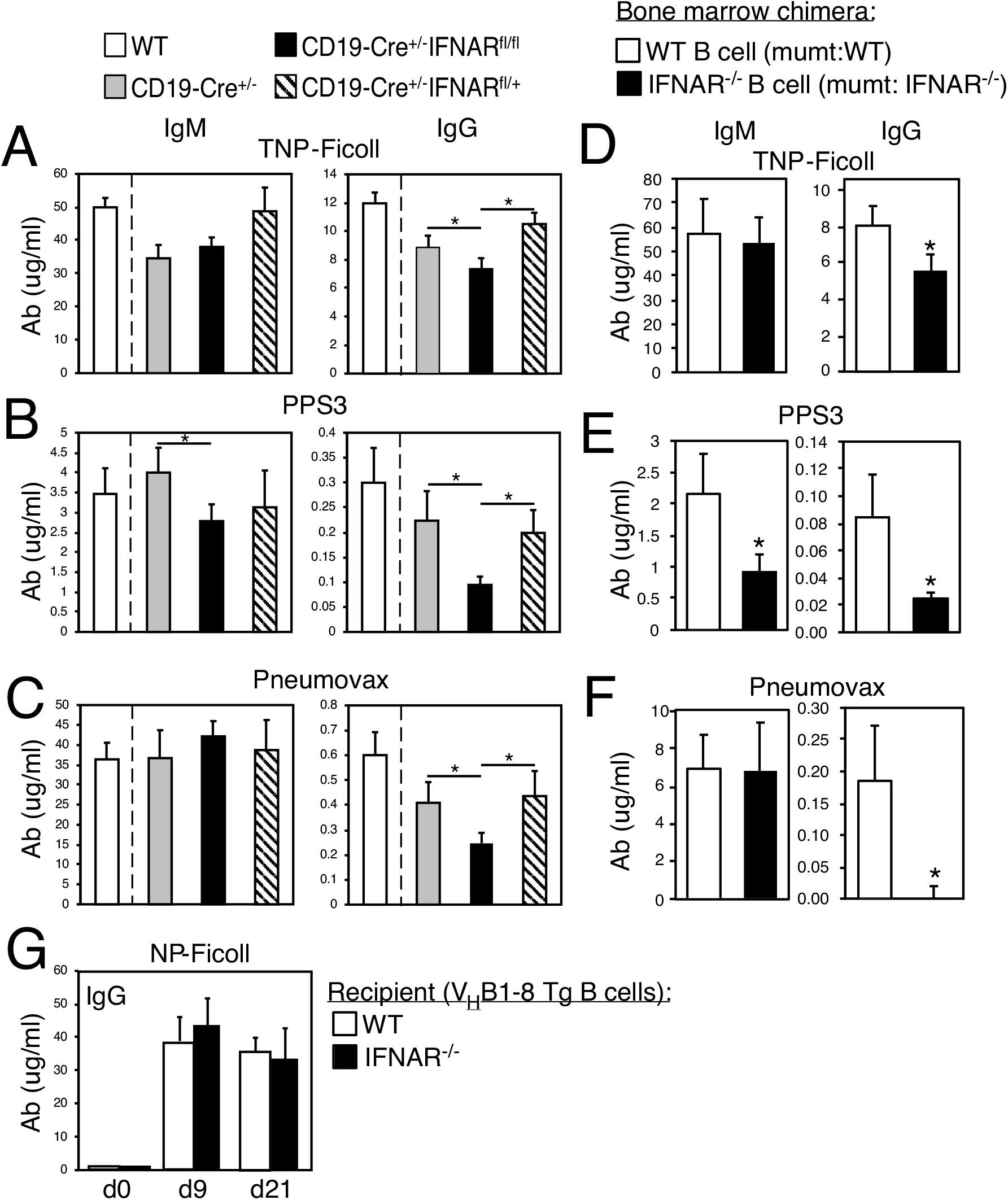
Type I IFN receptor signaling on B cells is required for optimal Ab responses to TI-2 Ags. A-C) WT, CD19-Cre^+/-^, CD19-Cre^+/-^FNAR1^fl/fl,^ and CD19-Cre^+/-^IFNAR1^fl/fl^ mice were immunized as in Figure 1, with serum IgM and IgG levels against TNP (A), PPS3 (B), and total Pneumovax (C) examined on day 21. Asterisks (*) indicate differences in values as compared to CD19-Cre^+/-^ mice. D-F) Irradiated WT mice reconstituted with mixed bone marrow from WT and mumt mice (10:90) or IFNAR^−/−^ and mumt mice (10:90) were rested for 8 weeks and then immunized as in A-C, with TNP (D), PPS3 (E), and Pneumovax-specific IgM and IgG assessed on day 27 post immunization (n=7/group). G) WT and IFNAR^−/−^ mice received 5 x 10^6^ V_H_B1-8 Tg spleen B cells i.v. and 5 x10^5^ peritoneal cavity B cells i.p. Recipients were immunized with 25 μg NP-Ficoll i.p. one day later and NP-specific IgG levels were assessed (n=4-5 mice/group). Results representative of 2 experiments.

We complemented the above studies by generating radiation bone marrow chimeras. These chimeras were constructed by reconstituting irradiated WT mice with bone marrow from B cell-deficient mumt mice mixed with either bone marrow from WT mice or IFNAR^−/−^ mice (90:10 ratio). As shown in Figure 3D, chimeras with B cells lacking IFNAR produced significantly less IgG (30% reduced) in response to TNP-Ficoll than chimeras reconstituted with WT B cells. IgM and IgG responses to PPS3 were also decreased 3 to 4-fold in chimeras with IFNAR-deficient B cells and in Pneumovax-specific ELISAs, IgG was barely detectable in these chimeras (Fig. 3E-F). Thus, mice with selective IFNAR deficiency on B cells exhibit impaired IgM and IgG responses to TI-2 Ags to a similar degree in mice lacking IFNAR on all cells. Finally, reconstitution of IFNAR^−/−^ mice with B cells from IFNAR^+/+^ V_H_B1-8 Tg mice in which a fraction of B cells (5-10%) coexpress the λ1 chain with the V_H_B1-8 transgene to yield a high affinity receptor specific for NP (11) yielded NP-specific IgG responses that were similar to WT recipients (Fig. 3G), indicating loss of non-B cell IFNAR expression did not noticeably impact this TI-2 Ab response. Thus, B cell-intrinsic IFNAR expression plays a major role in promoting TI-2 Ab responses.

### Type I IFN signaling supports early activation and expansion of Ag-specific B cells in response to TI-2 Ags

We assessed the early B cell response to TNP-Ficoll in WT and IFNAR^−/−^ mice to determine if defects in activation, expansion, and/or differentiation could be identified. As shown in Figure 4A, naïve WT and IFNAR^−/−^ mice had similar frequencies of TNP-specific CD19^+^ B cells in the spleen. However, 5 days following immunization, TNP-specific CD19^+^ B cell frequencies expanded 3-fold in WT mice, but only 2-fold in IFNAR^−/−^ mice (Fig. 4A). We noted similar decreases (~30%) in the frequencies of TNP-specific CD138^+^ and IgG^+^CD138^+^ plasmablasts as well as the frequencies of dividing (Ki67^+^) TNP-specific B cells, IgG^+^ B cells, and CD138^+^ B cells (Fig. 4A). The comparable reduction among total, CD138^+^, IgG^+^ and CD138^+^IgG^+^ cell populations in IFNAR^−/−^ mice, despite similar frequencies of naïve TNP-specific B cells suggested reduced TI-2 Ab responses in IFNAR^−/−^ mice may be due to a defect in early activation or expansion. Consistent with this, we found both TNP-specific CD138^neg^ B cells and differentiating CD138^+^ TNP-specific B cell plasmablasts in IFNAR^−/−^ mice had significantly lower levels of CD86 expression following immunization (Fig. 4B). Furthermore, CD11b expression on TNP-specific B cells from immunized IFNAR^−/−^ mice was significantly reduced, potentially indicative of impaired activation of CD11b-expressing B cells (ie., B-1b cells).

**Figure 4.**
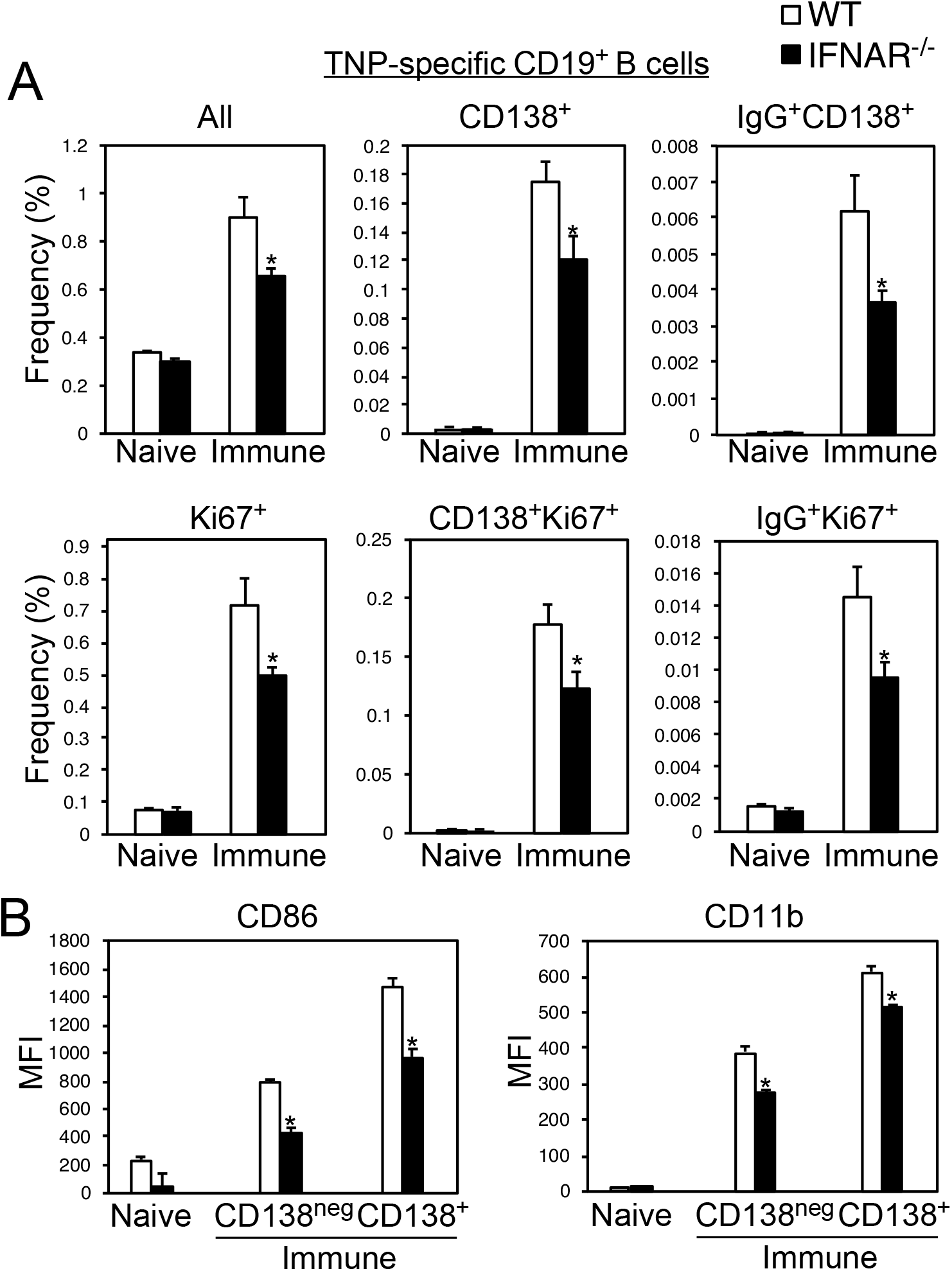
Impaired activation and expansion of TNP-specific B cells in IFNAR1^−/−^ mice following TNP-Ficoll immunization. (A-B) TNP-specific CD19^+^ spleen B cells were examined in naïve (n=4-5/group) and TNP-Ficoll-immunized (d5) WT and IFNAR1^−/−^ mice (n=5/group). The mean frequencies of TNP-specific CD19^+^, CD19^+^CD138^+^, CD19^+^IgG^+^CD138^+^, CD19^+^Ki67^+^, CD19^+^CD138^+^Ki67^+^, and CD19^+^IgG^+^Ki67^+^ cells among total splenocytes are indicated. B) Mean fluorescence intensity (MFI) of CD86 and CD11b among naïve and immune CD138^neg^ and CD138^+^ TNP-specific B cells in WT and IFNAR^−/−^ mice. Asterisks (*) indicate significant differences between values for WT and IFNAR^−/−^ mice.

### Type I IFN signaling is not required for MPL/TDCM adjuvant effects on Ab responses to TI-2 Ags

MPL/TDCM induces type I IFN production in vivo and does not optimally enhance TI-2 Ab responses in mice lacking the TIR-domain-containing adapter-inducing interferon-β (TRIF) adapter, which is an important inducer of IFN-β (8, 13). This, along with results from a previous study demonstrating type I IFN-mediated signaling on B cells was critical for poly(I:C)-mediated adjuvant effects in increasing IgG responses to NP-Ficoll (9) raised the possibility that type I IFN signaling was also critical for MPL/TDCM adjuvanticity. Given that IFNAR deficiency resulted in a more significant impairment of Ab responses to PPS relative to haptenated Ficoll and previous work showing that MPL/TDCM has more significant effects on increasing Ab responses to PPS relative to haptenated Ficoll (6), we assessed MPL/TDCM effects on responses of IFNAR^−/−^ mice to Pneumovax23 delivered i.m. As shown in Fig. 5, IFNAR^−/−^ mice exhibited significantly impaired PPS3- and Pneumovax23-specific IgM and IgG responses to Pneumovax23 delivered i.m., similar to results obtained for i.p. immunization (Fig. 1). However, inclusion of MPL/TDCM yielded significantly increased primary and secondary IgM and IgG responses to PPS3 and Pneumovax23, although IgG responses were variable. Thus, IFNAR is not required for MPL/TDCM adjuvant effects on PPS-specific Ab responses; moreover, MPL/TDCM overcomes defective Ab responses to Pneumovax23 associated with IFNAR deficiency.

**Figure 5.**
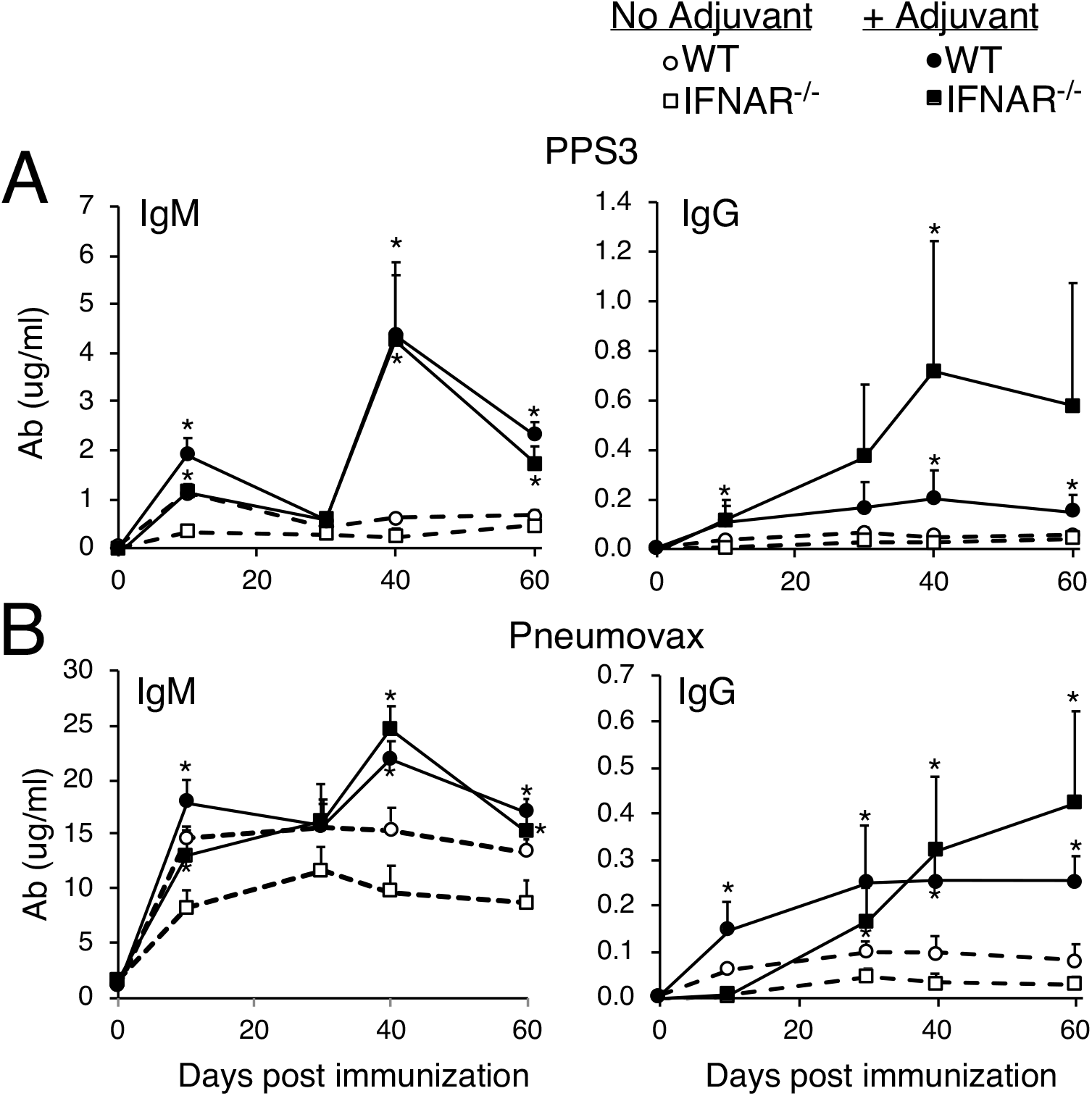
MPL/TDCM adjuvant effects on TI-2 Ab responses do not require type I IFN receptor signaling. A-B) WT and IFNAR1^−/−^ mice were immunized i.m. with Pneumovax23 containing 1 μg each PPS either alone or mixed with MPL/TDCM. Serum IgM and IgG levels reactive against PPS3 (A) and whole Pneumovax23 (B) were determined by ELISA. Asterisks (*) indicate significant differences between Ab levels in mice of the same genotype immunized with and without adjuvant (n=5 mice/group).

### B-1b and CD23^−^ spleen B cells produce increased Ab responses to NP-Ficoll coadministered with adjuvant, regardless of immunization route

Given the above findings and previous work demonstrating that a type I IFN-activating adjuvant elicited its IFNAR-dependent effects on Ab responses to NP-Ficoll through activating follicular B cells (9), we questioned whether MPL/TDCM enhanced TI-2 Ab responses by supporting Ab production by distinct B cell subsets. We found splenic and peritoneal NP-specific V_H_B1-8 Tg B cells, particularly CD11b^+^ peritoneal B-1b cells, were responsive to MPL/TDCM-supported Ab secreting cell (ASC) differentiation in the context of TI-2 Ag activation in vitro as evidenced by increased BLIMP1 expression by these cells in the presence of adjuvant (Supplemental Fig. 1A-B). To determine whether MPL/TDCM elicited its effects through select subsets modulated these effects, we performed adoptive transfer experiments using B cells from V_H_B1-8 Tg mice (11).

Naïve CD43^−^ peritoneal B-1b cells and CD23^+^ (enriched for follicular B cells) and CD23^−^ spleen B cells (enriched for marginal zone B cells) from V_H_B1-8 Tg mice were adoptively transferred i.v. into WT recipients which were then immunized with NP-Ficoll (i.p.). Importantly, λ1-expressing B cells are similarly represented among these subsets (11). B-1b cells produced the highest level of IgM and IgG in response to NP-Ficoll (Fig. 6A). Inclusion of MPL/TDCM also significantly increased NP-specific IgM^a^ (produced by Tg cells) and IgG (5–fold) by B-1b and CD23^−^ spleen B cells. However, MPL/TDCM had no effect on Ab production by CD23^+^ spleen cells.

**Figure 6.**
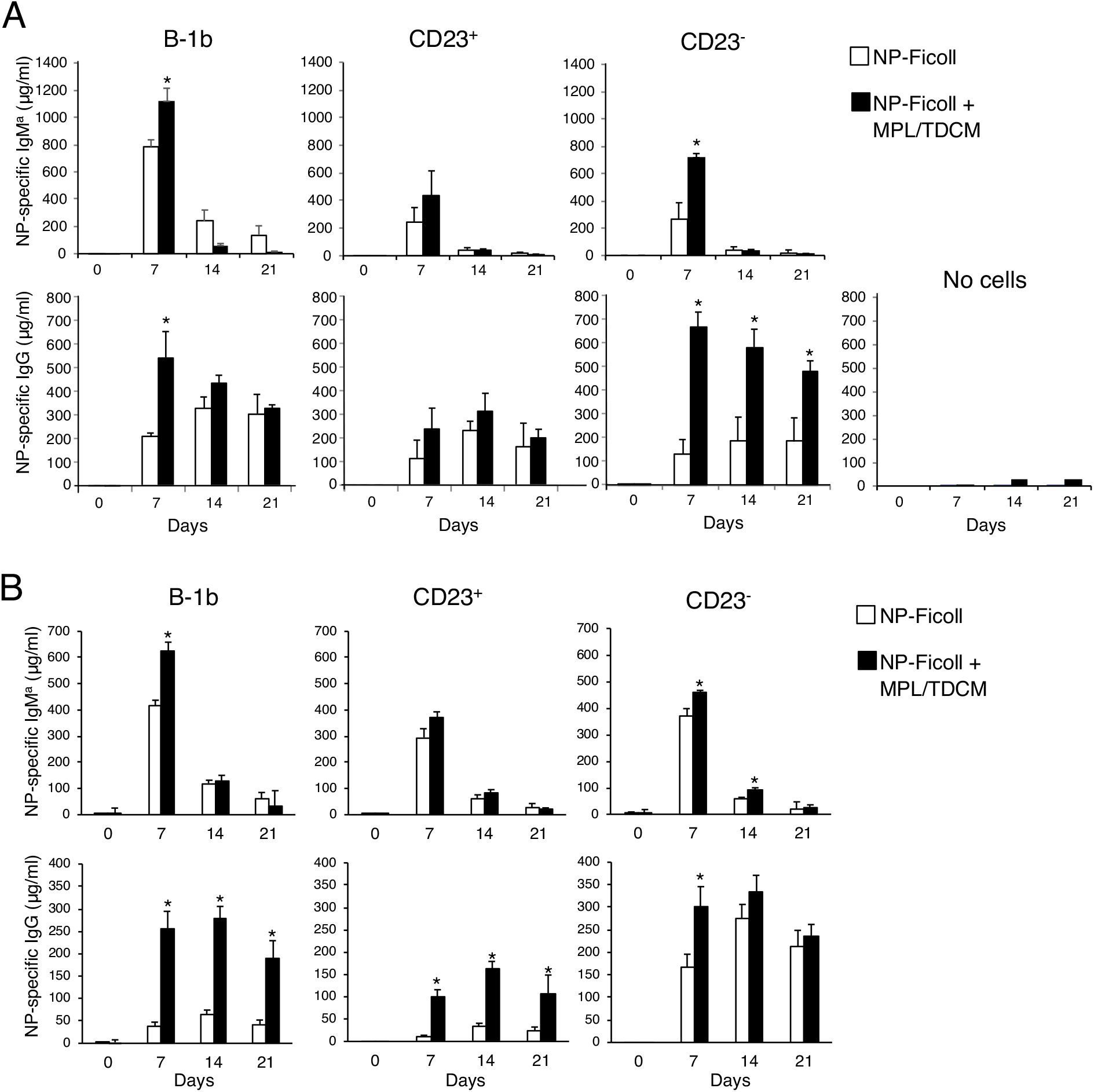
B-1b and CD23^−^ B cells produce increased Ab to NP-Ficoll coadministered with MPL/TDCM regardless of route. CD43^−^ V_H_B1-8 Tg CD11b^+^ peritoneal (B-1b) B cells (1.25 x 10^5^ in A, and 5 x 10^5^ in B), splenic CD23^+^ (5 x 10^5^) and splenic CD23^−^ (5 x 10^5^) B cells were transferred i.v. into WT recipients (n=4-5/group). One day later, recipients were immunized with 1 μg NP-Ficoll i.p. (A) or i.m. (B). NP-specific serum IgM^a^ and IgG levels are shown along with NP-specific IgG levels for immunized mice that did not receive V_H_B1-8 Tg cells (“no cells”). Results are representative of those obtained in two independent transfer experiments. Asterisks (*) indicate significant differences between Ab levels in recipients with and without adjuvant.

We next assessed whether i.m. immunization would impact Ab responses by distinct subsets. In response to NP-Ficoll delivered i.m., B-1b cells produced the highest amount of NP-specific IgM^a^, whereas CD23^−^ B cells produced the most IgG (Fig. 6B). MPL/TDCM significantly increased NP-specific IgM^a^ production from B-1b cells (1.5-fold) over recipient mice given adjuvant alone. Increases in IgM^a^ in recipients of CD23^−^ and CD23^+^ cells were moderate. MPL/TDCM increased NP-specific IgG production in recipients of each cell type, with d7 levels increased over 6-fold in B-1b and 9-fold in CD23^+^ recipients, but less than 2-fold in CD23^−^ B cell recipients. Thus, in the NP-Ficoll immunization system using high affinity V_H_B1-8 Tg B cells, B-1b and CD23^−^ spleen cells preferentially differentiate into ASC regardless of immunization route and MPL/TDCM further potentiates their level of Ab production, although there is nonetheless some responsiveness to MPL/TDCM in the V_H_B1-8 Tg CD23^+^ B cell population following i.m. immunization.

### B-1b and CD23^−^ spleen B cells produce increased Ab responses to Pneumovax23 coadministered with adjuvant

We next examined effects of MPL/TDCM on responses of B cell subpopulations to the pneumococcal vaccine, Pneumovax23. In muMT mice reconstituted with high numbers of spleen cells (3×10^7^), IgM and IgG levels against Pneumovax23 as well as serotype 3 PPS (PPS3) were not significantly different among recipients that received no immunization, Pneumovax only, or MPL/TDCM only (i.p.) whereas mice receiving Pneumovax plus MPL/TDCM i.p. produced significantly increased Pneumovax-specific IgM and IgG (Fig. 7A). PPS3-specific IgM and IgG was also increased, although not significantly. Thus, there is PPS-specific spleen B cell population that responds to MPL/TDCM adjuvant effects.

**Figure 7.**
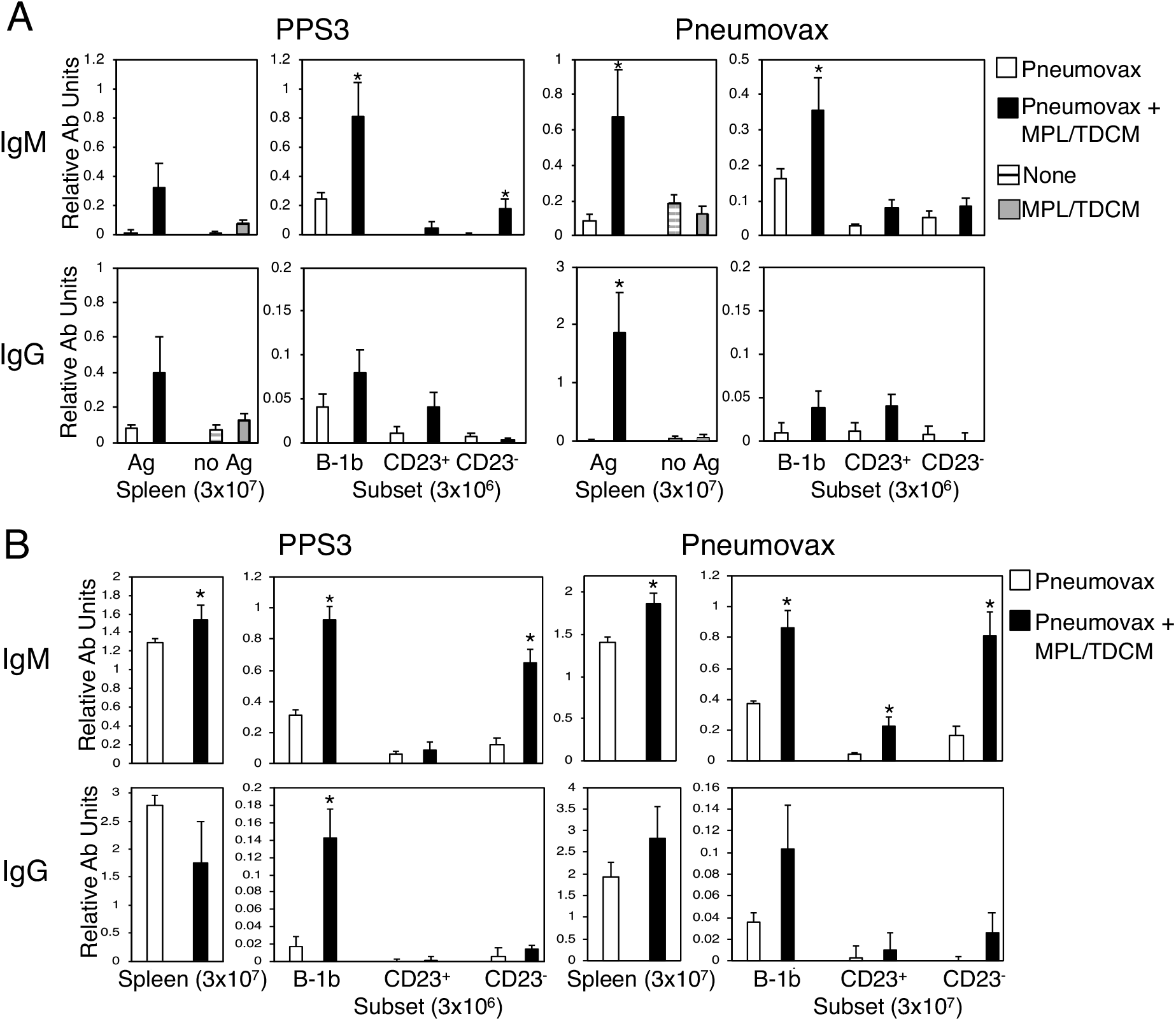
MPL/TDCM promotes increased B-1b and splenic CD23^−^ Ab responses to PPS, regardless of immunization route. WT splenocytes (3 x10^7^) or enriched peritoneal B-1b B cells, splenic CD23^+^ and CD23^−^ B cells (3 x 10^6^) were transferred into muMT recipients (n=4-7/group). One day later, recipients were immunized with Pneumovax23 either alone or with MPL/TDCM i.p. (A) or i.m.(B). An additional group of splenocyte recipients received no immunization (horizontal gray stripe) or MPL/TDCM only (gray fill) in (A). IgM and IgG recognizing PPS3 and total Pneumovax23 were assessed on d10 post immunization. Asterisks (*) indicate significant differences between Ab levels in recipients with and without adjuvant.

When examining sorted B cell populations, we found recipients of B-1b cells produced the highest level of Pneumovax23- and PPS3-specific IgM and PPS3-specific IgG in response to Pneumovax23 i.p. (Fig. 7A). MPL/TDCM significantly increased total Pneumovax23-specific IgM in B-1b, but not in CD23^−^ and CD23^+^ B cell recipients. Similarly, MPL/TDCM significantly increased PPS3-specific IgM production in B-1b cell recipients, and to a lesser extent in CD23^−^, but not CD23^+^ B cell recipients. PPS3- and Pneumovax-specific IgG was low but MPL/TDCM produced moderate increases in B-1b recipients.

In response to Pneumovax23 plus adjuvant delivered i.m., recipients of high dose splenocytes produced increased Pneumovax23- and PPS3-specific IgM, but not IgG (Fig. 7B). Recipients of B-1b cells and splenic CD23^−^ B cells produced significantly increased PPS3- and Pneumovax23-specific IgM in response to adjuvant (Fig. 7B). MPL/TDCM significantly increased PPS3-specific IgG as well as increased Pneumovax23-specific IgG, only in B-1b cell recipients. CD23^+^ B cell recipients were not very responsive. Thus, MPL/TDCM adjuvant effects observed in the context of Pneumovax23 immunization are largely carried out through innate B cell populations.

## Discussion

Antibody responses to polysaccharide Ags are critical for protection against encapsulated pathogens. However, our understanding of the factors regulating these responses remains incomplete. Our currrent study reveals several novel findings regarding regulation of TI-2 Ab responses. First, we demonstrate type I IFN signaling on B cells supports optimal Ab responses to TI-2 Ags, including pneumococcal polysaccharides, via support of early B cell activation and expansion. Second, we demonstrate an MPL-based adjuvant that significantly augments primary and secondary responses to TI-2 Ags (6), induces type I IFN production in vivo (7), and requires TRIF signaling for optimal adjuvant effects (6), does not require type I IFN for its effects and moreover, restores defective TI-2 Ab responses caused by IFNAR deficiency. Finally, we demonstrate MPL/TDCM elicits its adjuvant effects by largely promoting Ab production by innate-like B cells (ie., B-1b and CD23^−^ B cells). Collectively, our results provide valuable information regarding the role of type I IFN in supporting polysaccharide-specific Ab responses as well as critical information that may be leveraged for designing strategies to improve vaccines targeting polysaccharide Ags in the future.

The role of IFNAR in regulating B cell responses is complex based on findings reported for IFNAR^−/−^ mice and in vitro studies of B cells. We show IFNAR deficiency on a C57BL/6 background does not impact peritoneal or splenic B cell subset frequencies or numbers. An earlier study reported IFNAR^−/−^ mice on a 129sv background had normal absolute B cell numbers in bone marrow and spleen; however, it indicated the repertoire of immature bone marrow B cells was altered, and further, showed IFN I could modulate the sensitivity of activated B cells to BCR-dependent inhibition of terminal differentiation resulting from LPS or CD40 + IL 4 stimulation (14). In contrast, another study reported that IFN I in conjunction with IL-6 promotes plasma cell differentiation (15). Consistent with the former study, we found that addition of type I IFN to V_H_B1-8 Tg cells activated via TI-2 Ag stimulation in vitro (ie, NP-Ficoll, IL-5, and IFN-γ; (16)) resulted in reduced recovery of CD138^+^ cells (unpublished observations). Nonetheless, IFN I protects B cells against apoptosis and has been reported to enhance BCR-mediated activation and proliferation in some studies but have the opposite effect in others (17–20). Thus, both stimulatory and inhibitory effects of IFN-αβ on in vitro B cell proliferation and Ig production have been reported, perhaps due to the fact that high dose IFN I is inhibitory whereas low dose IFN is stimulatory for Ig production by activated B cells (21). Its positive role in supporting TNP-Ficoll-specific B cell activation and expansion in vivo in the absence of strong type I IFN production is consistent with this notion.

Our results demonstrate a clear role for B cell-expressed IFNAR in supporting TI-2 Ab production and reveals both IgM and IgG responses to PPS within the pneumococcal vaccine are critically dependent on type I IFN signaling. Work on the effects of type I IFN on TI Ab responses has been limited and largely restricted to anti-viral responses. The effect of type I IFN on B cell responses to vesicular stomatitis virus (VSV), which displays highly ordered VSV-G and induces TI IgM responses has been examined in two studies. One study demonstrated IFN-a enhanced VSV-specific B cell IgM production in response to UV-inactivated VSV in vitro and showed that B cell-expressed IFNAR was required for optimal IgM, but not IgG responses, to live VSV in vivo (22). In contrast, in response to noninfective VSV-viral like particles (VLP), IFNAR deficiency significantly reduced TD IgG responses but only slightly reduced IgM responses. This regulation was through B cell-extrinsic IFNAR expression, and the study reported little effect of B cell-expressed IFNAR on the TI Ab response to VSV-VLP (23). Finally, with regard to innate B1 cell production of Abs, IFNAR is required for B1 cell lymph node trafficking and IgM production in response to influenza virus infection due to its role in regulating CD11b conformational changes (24). However, it is not required for monophosphoryl lipid A-induced tumor reactive natural IgM production by B1a cells (7), despite its dependence on the TLR4-TRIF activation pathway (25). Given the significantly reduced activation and expansion of IFNAR^−/−^ TNP-specific B cells 5 days post TNP-Ficoll immunization, along with early effects of IFNAR deficiency on both IgM and IgG production, IFNAR signaling appears to be important for supporting early B cell activation events required for optimal responses to TI-2 Ags. Future work will determine the extent to which decreased CD11b expression by responding IFNAR^−/−^ B cells may be due to impaired activation of Ag-specific B cells and the extent to which reduced B-1b cell recruitment/activation is involved. Although IFNAR1 deficiency yields a distinct phenotype in humans relative to mice with regard to mucosal viral infections (26), whether mouse and human B cells share similar dependency on type I IFN for optimal TI-2 Ab responses remains unknown. Recent work defining the effects of mutations involved in altered type I IFN pathways along with analysis of patients either treated with IFN therapeutics or who have developed autoAbs against IFNs (26–29) will likely shed light on this in the future.

We investigated the role type I IFN plays in the adjuvant effects of MPL/TDCM on TI-2 Ab responses in the current study due to: 1) its dependency on TRIF signaling (6), 2) results from a previous study demonstrating the requirement for B cell-expressed IFNAR in the adjuvant effects of poly(I:C) on TI-2 Ab responses (9), and 3) the dependency of non-adjuvanted TI-2 Ab responses on B cell-expressed IFNAR identified herein. Nonetheless, we found MPL/TDCM works independently of type I IFN signaling and largely overcomes defective TI-2 Ab responses in IFNAR^−/−^ mice. The TLR4-dependent effects of MPL on activating both MyD88 and TRIF pathways in B cells (6), the latter of which activates NF-kB in addition to type I IFN production, may explain these findings.

Our work demonstrates MPL/TDCM significantly enhances Ab responses to polysaccharide Ag largely through its effects on innate-like B cells (ie., B-1b and CD23^−^ spleen B cells). The adjuvant effects on increasing PPS-specific IgG production by B-1b cells to Pneumovax23 delivered i.m. was particularly notable. B-1 and MZ B cells are most often found producing Abs against TI-2 Ags in the absence of adjuvant, so it is perhaps not surprising that this adjuvant further promotes the ability of these B cells to produce more IgM and in some cases IgG, against these polysaccharide Ags (11, 30). Although V_H_B-1-8 Tg CD23^+^ spleen B cells showed some responsiveness to adjuvant with NP-Ficoll delivered i.m., this population responded poorly when PPS was used. These findings contrast with a previous study that showed follicular B cells produce increased IgG responses to NP-Ficoll delivered i.p. when poly(I:C) is used as an adjuvant. This was shown to depend on the costimulatory effects of type I IFN on B cells as opposed to direct TLR3 signaling (9). In addition to being less responsive to TI-2 Ags relative to marginal zone and B-1 cells, follicular B cells are also less responsive to TLR agonists (31). In contrast to the mechanism defined for poly(I:C), the MPL-based adjuvant functions through direct activation of TLR4 on B cells (6). This supports the possibility that combinations of distinct pattern recognition receptor agonists that work through different pathways may be optimized to significantly improve B cell responses to polysaccharide Ags in humans in the future.

## Notes

This work was supported by NIAID/NIH R01AI18876 and R21AI144758. MAS was supported by NIH T32AI007401. Research reported in this publication was also supported by the National Center for Advancing Translational Sciences of the National Institutes of Health under Award Number UL1TR001420. The content is solely the responsibility of the authors and does not necessarily represent the official views of the National Institutes of Health. The authors have no conflicts of interest to disclose.

### Competing Interest Statement

The authors have declared no competing interest.

