## Supplemental Figure 1 for "Type 1 interferon supports B cell responses to polysaccharide antigens but is not required for MPL/TDCM adjuvant effects on innate B cells"

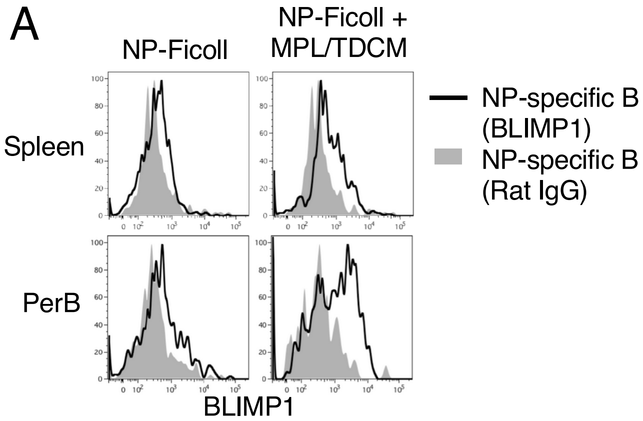


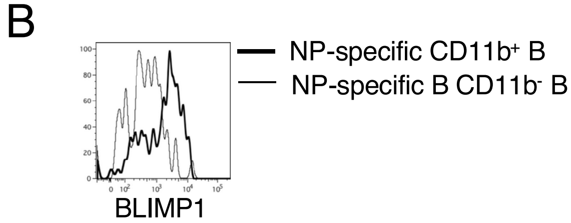


**Supplemental Figure 1. MPL/TDCM promotes increased upregulation of BLIMP1 on NP-specific spleen and peritoneal B cells activated with NP-Ficoll**

**A-B)** $V_H$B1-8 Tg NP-specific splenocytes and peritoneal cells were cultured at 1 x $10^6$ cells/ml in complete RPMI containing 10% FCS in 96-well culture plates in triplicate and stimulated with $NP_{40}$-Ficoll (100 ng/ml) alone or with MPL/TDCM adjuvant (Sigma) containing 1 μg/ml MPL and TDCM. On day 2, NP-binding B220$^+$ B cells (A) and B220$^+$CD11b$^+$ and CD11b$^-$ peritoneal B cells (B) were analyzed for intracellular BLIMP1 expression by flow cytometry using eBioscience FoxP3/Transcription factor staining buffer set in conjunction with BLIMP1 (clone 5E7) staining. Rat IgG2b was substituted for BLIMP1 during intracellular staining to determine background level staining. Cells were acquired using a FortessaX20 cytometer (BD Biosciences) with FSC-A/FSC-H doublet exclusion. Data were analyzed using FlowJo analysis software (Tree Star). Gray shaded histograms show binding for PE-labeled control rat IgG1 which was substituted for anti-BLIMP1 PE in the staining panel.
